# Dissecting cell-free DNA fragmentation variation in tumors using cell line-derived xenograft mouse

**DOI:** 10.1101/2024.07.03.601978

**Authors:** Ruiqing Fu, He Amy Su, Yafei Tian, Hongyan Chen, Daru Lu

## Abstract

Cell-free DNA (cfDNA) is increasingly studied for its diverse applications in non-invasive detection. Non-randomly cleaved by nucleases and released into the bloodstream, cfDNA exhibits a variety of intrinsic fragmentation patterns indicative of cell status. Particularly, these fragmentation patterns have recently been demonstrated to be effective in predicting cancer and its tissue-of-origin, owing to increased variation of fragmentation features observed in tumor patients. However, there remains a lack of detailed exploration of altered cfDNA fragmentation profiles in tumors, which consist of a mixture of both non-tumor cfDNA and circulating tumor DNA (ctDNA). Hence, we leveraged the human tumor cell line-derived xenograft (CDX) mouse model, where different tumor cell lines were implanted into different anatomical sites, to isolate pure ctDNA and separately investigate the fragment properties of CDX-induced cfDNA and ctDNA. We found an enrichment of short cfDNA fragments in both CDX-induced cfDNA and ctDNA compared to normal plasma cfDNA, with more elevated short fragments in ctDNA. Moreover, the CDX-induced cfDNA fragmentation features distinguished between CDX models of different tumor cell lines, while the ctDNA fragmentation features conversely discriminate between CDX models of different anatomical sites. The results suggested that both non-tumor cfDNA and ctDNA contribute to the increased variation observed in tumors, and that cfDNA fragmentation may be highly variable and susceptible to regulations by both original cells and cells within the local niche.

## Introduction

Liquid biopsies, particularly the blood-based tests, are recently actively developed, for its promising applications in clinical oncology [1-3], prenatal testing [4, 5], and organ transplant monitoring [6, 7]. Among a range of blood biomarkers, the plasma cell-free DNA (cfDNA) is widely studied and applied, especially for early cancer detection [8-11], owing to its non-invasive accessibility, and rich molecular characteristics, including genomic mutations [12, 13], methylation alterations [14, 15], and fragmentomics signatures [16, 17], which could inform cell pathological status (*e*.*g*., cancer cells). The genomic mutations and methylation alterations of cfDNA have been intensively investigated due to their roles in tumorigenesis [18, 19] and cancer prediction, while fragmentomics is a novel feature that encompasses various properties of cfDNA fragments, including their size, end motifs and orientation, nucleosome occupancy, and topology [20-22].

The cfDNA molecule typically wraps around a nucleosome core, corresponding to ∼147 bp, with an unbound linker (∼20 bp) interspersed between nucleosomes [23]. During various cell death processes [24], cfDNA undergoes non-random cleavage by a series of nucleases [25], where the DNA is naked, and is released into the bloodstream, thus resulting in a distinctive distribution of fragment size, with a mode of ∼167 bp [26], and preferred frequencies of fragment end motifs [27]. Meanwhile, nucleosomes that wrapped by DNA and transcription factors that bind to DNA can protect it from cleavage, imprinting the nucleosome occupancy [26]. Similarly, other factors that modify DNA conformation, such as DNA methylation, also play a role in cfDNA fragmentation [28]. In other words, cfDNA fragments from open regions or regions depleted of other modification factors may be cleaved too shortly to be captured for sequencing by conventional sequencing approaches, thus resulting in very low genome coverage [17, 29]. The majority of cfDNA originates from blood cells [5, 26], while in cancer patients, it contains circulating tumor DNA (ctDNA) derived from tumor cells [30]. As a result, fragmentomics signatures can be utilized to predict cancer, as well as tissue-of-origin in principle [26]. Studies have manifested an enrichment of shorter cfDNA fragments in the presence of tumors [31], accompanied by increased variation in fragment profiles among cancer patients [16]. Therefore, despite the alterations in genomic variation and methylation, the presence of tumors may also change the composition of cfDNA fragments. However, it remains to be elucidated that whether such tumor-derived variation is solely contributed by ctDNA or by both ctDNA and non-tumor cfDNA. Several studies have attempted to infer tissue-of-origin of cancers using fragmentomics signatures [32-34], but these efforts have been limited by small sample sizes and specific cancer types. There has been a lack of detailed exploration into using cfDNA fragmentation to differentiate among cell types. The natural mixture of cfDNA fragments and the low concentrations of ctDNA pose significant challenges in dissecting their properties. Thus, we leveraged cell line-derived xenograft (CDX) mouse models to isolate nearly pure ctDNA from cfDNA and separately investigated their fragmentation properties, demonstrating the influence of tumor cells on the non-tumor cfDNA, and their efficiency in differentiating cancer origins.

## Results

### Isolating ctDNA from CDX-induced cfDNA by CDX models

To separately and accurately characterize the properties of plasma cfDNA and ctDNA, we established 12 CDX mouse models, by implanting A549 lung carcinoma (A549) and MHCC-97H liver carcinoma (M97H) cell lines into both the pancreas and rectum of nude mice (n = 3 per group). The CDX models allowed us to delineate variations in cfDNA fragmentation attributed to tumor cell lines and anatomical sites of tumorigenesis (**Figure 1A**). Whole blood was collected from the CDX models for plasma cfDNA extraction, followed by deep whole genome sequencing (WGS) for each model (mean coverage >38×, with a range of 33.18-42.78×). Additionally, three normal mice were used as controls to provide the normal background of cfDNA profile. The extracted plasma cfDNA from the CDX models comprised a mixture of non-tumor cfDNA released from mouse cells (*i*.*e*., CDX-induced cfDNA) and tumor cfDNA derived from human tumor cell lines (*i*.*e*., ctDNA). Several tools have been developed to analyze xenograft sequencing data, with a variable of sensitivities to detect xenograft reads [35]. With the aim of minimizing interference between CDX-induced cfDNA and ctDNA, while maximizing the retrieval of human-derived reads, we built a bioinformatic pipeline to distinguish between the two sources of cfDNA (see **Methods**). Briefly, we first generated a list of black regions in the human genome, which were highly similar to the mouse genome, by mapping human-prone reads from control mouse samples to the human genome. Subsequently, sequencing reads identified as human genomic fragments by either xenome [36] or XenofilteR [37] were re-aligned to the human reference genome (hg19), after which high-quality alignments that did not fall within the black regions were then used for ctDNA analysis (**Figure S1**). With this elaborate subtraction process, we were able to isolate pure human-derived ctDNA.

**Figure 1.**
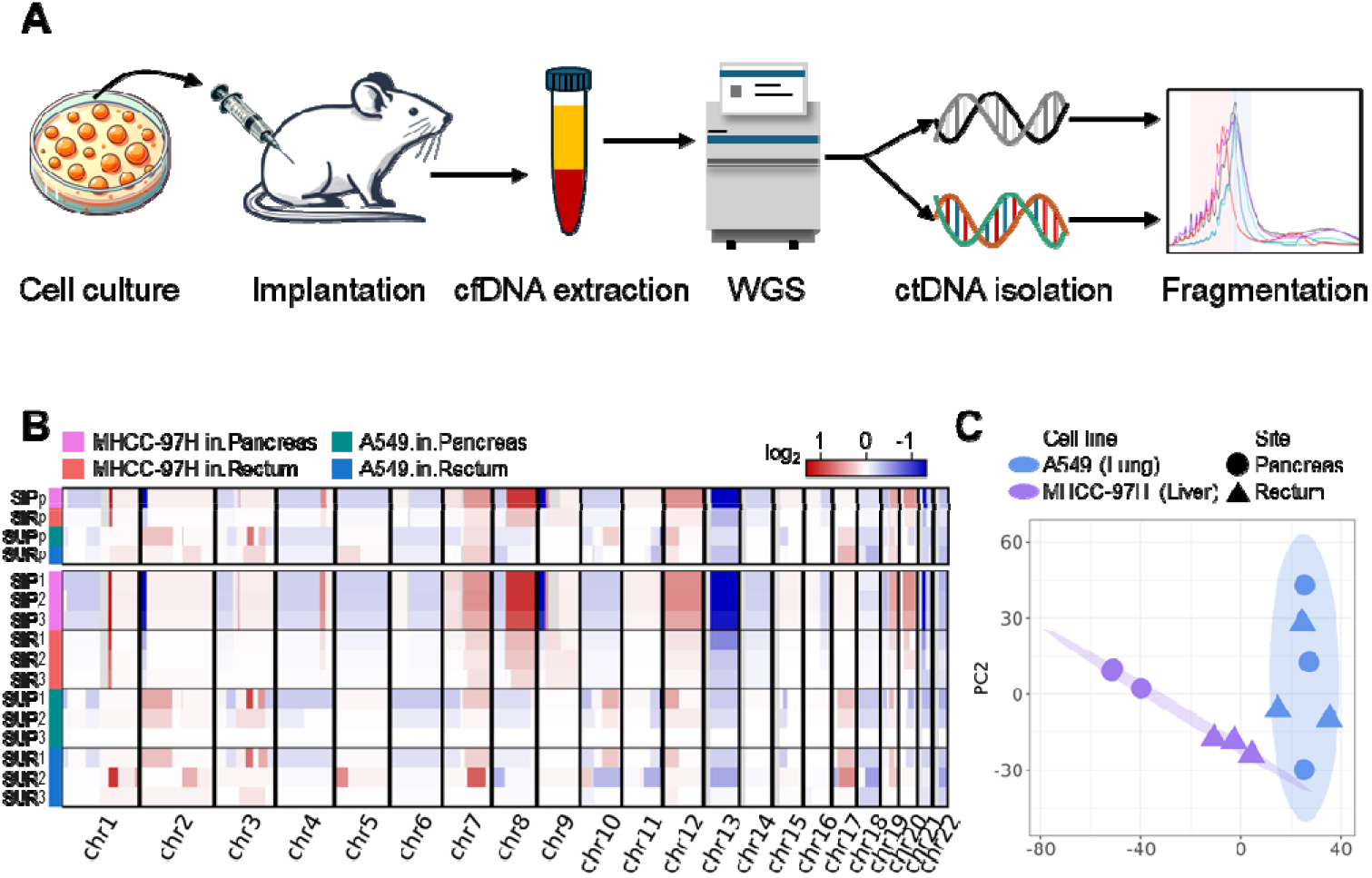
Schematic illustration of the study and CNA patterns in tumor cells. (A) The overview of the study design. (B) The CNAs detected by isolated ctDNA fragments in human-derived tumor cells from each CDX model (bottom panel) and in pooled representative samples (top panel). (C) Principal component analysis (PCA) on the CNA patterns for CDX models

The fractions of ctDNA ranged from 0.21% to 18.96% (**Table S1**), resulting in ctDNA fragments (50-1000 bp) ranging from 2.99×10^5^ to 424.57×10^5^, with the M97H cell in pancreas models exhibiting the highest proportion of ctDNA. To enhance the signal of ctDNA, we pooled the fragments together for each of the four groups. As expected, the ctDNA fragments manifested a diverse spectrum of copy number alterations (CNAs) across CDX models (**Figure 1B**), with the exception of SUP3, possibly due to its enrichment in very short fragments (**Figure S5**), which may not be suitable for large-scale CNA detection. Interestingly, despite substantial differences between the two tumor cell lines, we also observed considerable variations between the two anatomical site models within each implanted cell line. Based on the CNA patterns, the models were first clustered by different cell lines and then by different implantation sites (**Figure 1C**), suggesting potential differentiation of ctDNA fragments post-implantation, as CNAs were mainly inferred from the sequencing coverage of ctDNA fragments.

### Fragmentation variations revealed by CDX-induced cfDNA

To explore the properties of CDX-induced cfDNA potentially influenced by tumorigenesis, we first analyzed mouse cfDNA. All mouse cfDNA data were down-sampled to ∼200 million reads, corresponding to an average of 995.92×10^5^ and 993.87×10^5^ cfDNA fragments for normal plasma samples and xenograft plasma samples, respectively. With the protection of mono-nucleosomes, the majority of cfDNA fragments typically exhibited a size of ∼167bp [26], with periodic decreases of ∼10 bp (**Figure 2A** and **S2**). However, compared to normal plasma cfDNA, the CDX models exhibited an increased proportion of short cfDNA fragments. Therefore, we calculated the ratio between short fragments (80-160 bp) and long fragments (161-200 bp) to characterize the enrichment of short fragments of cfDNA, referred to as the S2L ratio. Interestingly, CDX models showed significantly increased short cfDNA fragments (p = 0.048 and 0.048 for A549 and M97H CDX models, respectively; U-test) (**Figure 2B**). Furthermore, the clustering of mouse cfDNA also revealed distinct fragment distributions between normal plasma samples and xenograft plasma samples, and even between the two CDX models implanted with different tumor cell lines (**Figure 2C** and **S3**). The differences between normal plasma cfDNA and xenograft plasma cfDNA (*i*.*e*., CDX-induced cfDNA) suggested alterations in cfDNA fragmentation in the microenvironment in the presence of tumor cells.

**Figure 2.**
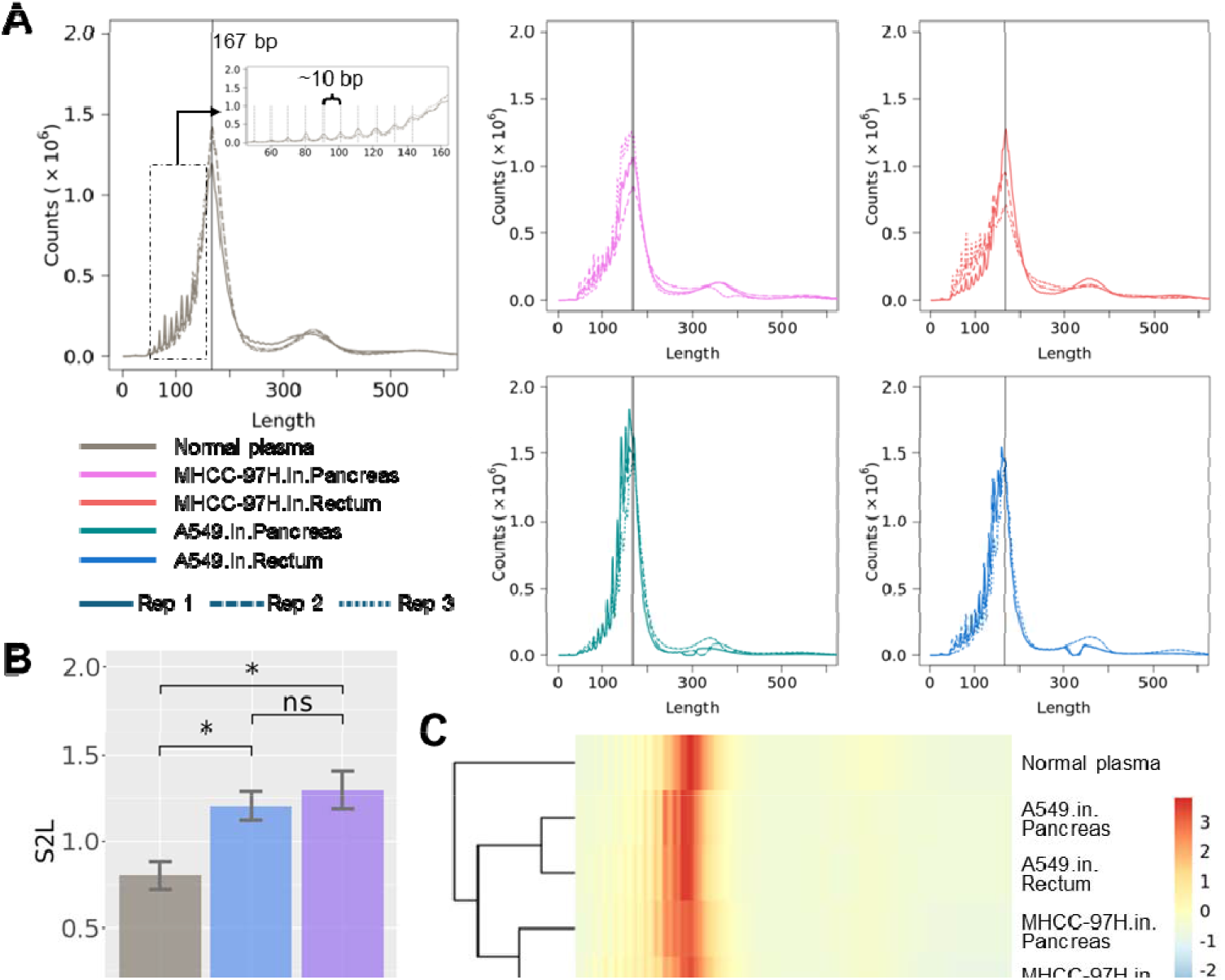
Fragment profile of mouse plasma cfDNA and the enrichment of short fragments. (A) The fragment size distribution of normal control plasma cfDNA, and CDX-induced cfDNA from four types of CDX models, with a major peak at 167 bp. (B) Enrichment of short fragments in CDX-induced cfDNA from CDX models, indicated by S2L ratio. (C) Hierarchical clustering of cfDNA fragment size in pooled samples. (significance: ‘*’, p < 0.05; U-test)

To further investigate the differences between CDX models based on different tumor cell lines and different anatomical sites, we calculated several fragmentomics features of the cfDNA fragments, including breakpoint motif (BPM), end motif (EDM), and fragment size distribution (FSD). Due to the limited sample size, no significant differences (p > 0.05; U-test) were observed neither between anatomical sites within each cell line, nor between cell lines within each anatomical site. However, we found that CDX models can generally be clustered by different cell lines, using these features (Figure 3A and S4). Therefore, we grouped the CDX models either by tumor cells or by anatomical sites and attempted to detect informative markers (*i*.*e*., motifs or FSD bins) between different groups by calculating the area under curve (AUC) values. Interestingly, we found that the distribution of marker AUCs between tumor cell lines were significantly different from that between anatomical sites (p < 1e^-4^; U-test) (Figure 3B), with a significantly greater number of informative motifs and FSD bins (AUC > 0.9) detected between cell lines compared to those between anatomical sites (p < 1e^-4^; χ2-test) (**Table 1**). The feature of fragment size, namely the FSD, exhibited an enhanced ability to distinguish between cell lines rather than anatomical sites. To further assess the significance of FSD in distinguishing samples, we performed permutations (n = 1,000) of the CDX models, resulting in a significant differentiation between two tumor cell lines (p = 0.03) but not between two anatomical sites (p = 0.57). The FSD feature consisted of 5bp-bins ranging from 65 bp to 399 bp and was calculated by counting the cfDNA fragments falling within the corresponding bin size for each individual chromosome (see **Methods**). Notably, we found that M97H cell CDX models exhibited enriched cfDNA fragments in the size range of 220-309 bp, while A549 cell CDX models involved more shorter cfDNA fragments ranging from 150 bp to 174 bp (**Figure 3C**). The results indicated that different tumor cells may result in microenvironments with considerable differences between each other.

**Figure 3.**
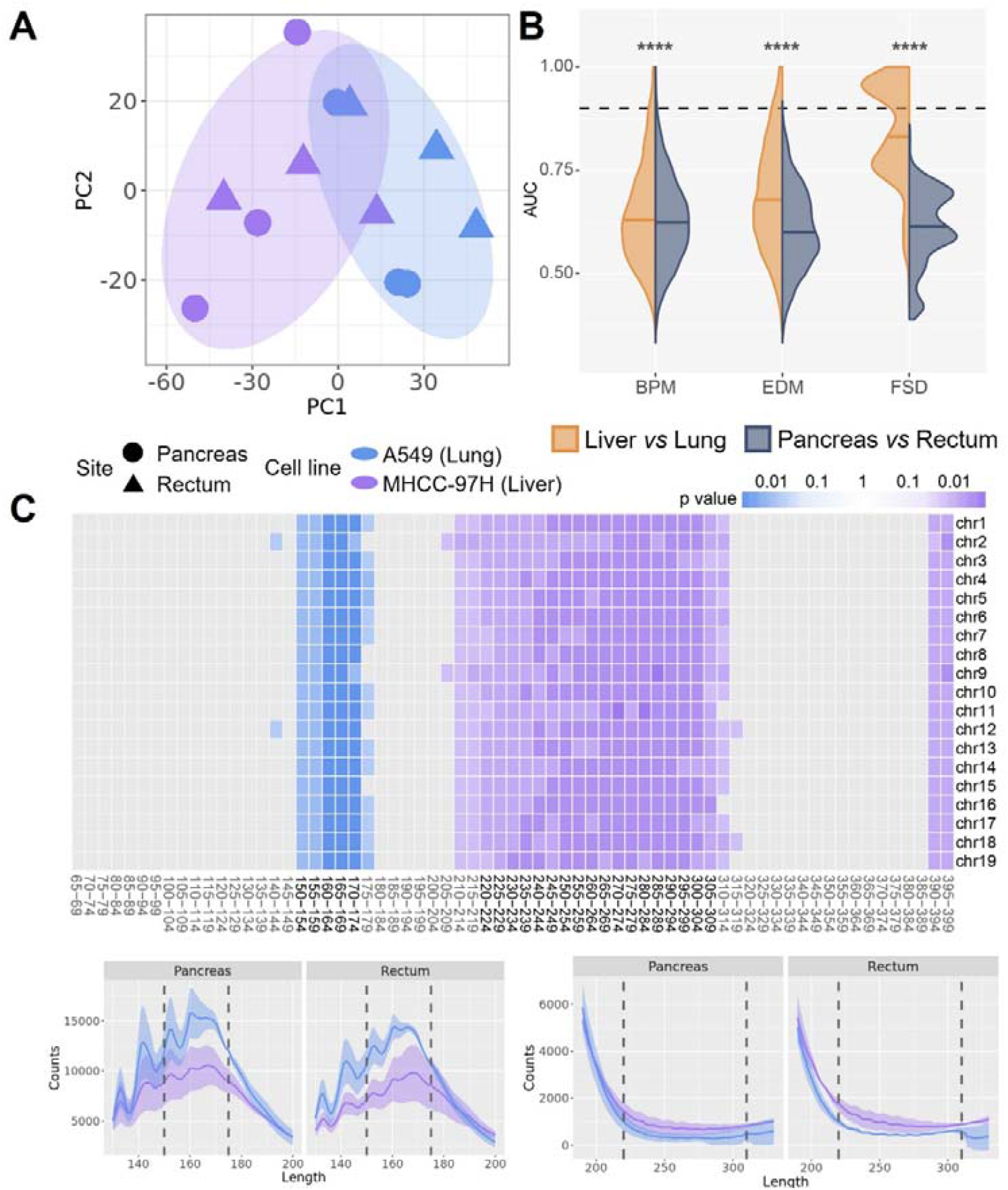
CDX-induced cfDNA distinguished between different cell line CDX models. (A) Different cell line CDX models are distinguished mainly by CDX-induced cfDNA fragmentation features like fragment size distribution (FSD). (B) Comparison of AUC value distributions between samples grouped by tumor cell line and by anatomical site, for fragmentation features including breakpoint motif (BPM), end motif (EDM), and FSD, respectively. The dashed line indicates AUC = 0.9.(significance: ‘****’, p < 1e^-4^; U-test) (C) The fragment bin sizes of FSD that are enriched in M97H CDX models (purple) and A549 CDX models (blue) (top panel), and the differentiation of the fragment profiles corresponding to the bin sizes, with the line indicates the mean value of the replicates and the shadow indicates the standard error (bottom panel). The mosaic was colored if the AUC of the corresponding FSD bin size was larger than 0.9, with the darkness indicating the p value calculated by the U-test.

**Table 1.** Number of informative fragmentation features (AUC > 0.9)

| AUC > 0.9 | cfDNA |  |  | ctDNA |  |  |
| --- | --- | --- | --- | --- | --- | --- |
| Feature | BPM | EDM | FSD | BPM | EDM | FSD |

**Table 1.** Number of informative fragmentation features (AUC > 0.9)
|  | (n=4096) | (n=4096) | (n=1407) | (n=4096) | (n=4096) | (n=2680) |
| --- | --- | --- | --- | --- | --- | --- |
| MHCC-97H vs. A549 | 163 | 257 | 589 | 55 | 71 | 4 |
| Pancreas vs. Rectum | 26 | 3 | 0 | 157 | 185 | 278 |

### Fragmentation variations revealed by human-derived ctDNA

We sought to investigate the potential differences between original tissues or cell types, possibly revealed by ctDNA fragments. Unlike cfDNA fragments, ctDNA was enriched in shorter fragments, with a peak at ∼143 bp (**Figure 4A** and **S5**), consistent with previous studies [31, 38]. The periodicity was also significantly shorter than that in cfDNA (7.17 bp and 9.98 bp; p = 0.021; paired t-test). The S2L ratio demonstrated that ctDNA involved a significantly higher proportion of short fragments than both normal plasma cfDNA (p = 0.024 and 0.024 for M97H cell and A549 cell CDX models, respectively; U-test) and CDX-induced cfDNA (p = 0.031 and 0.031 for M97H cell and A549 cell CDX models, respectively; paired U-test). Notably, there were no significant differences between CDX models of the two cell lines, but ctDNA from pancreas models was significantly more enriched in short fragments than that from rectum models (p = 0.0087; U-test) (**Figure 4B**). We further conducted clustering on the ctDNA from CDX models and found that samples from the same anatomical sites clustered together, rather than from the same cell line (**Figure 4C** and **S6**). The ctDNA fragments from pancreas models were enriched in a relatively narrow range of length, while the fragments from rectum models exhibited a much wider range of length. The SUP3 sample, which was not observed with detectable large-scale CNAs, displayed an outlier pattern characterized by the highly enriched presence of very short fragments (less than 90 bp) (**Figure S5**).

**Figure 4.**
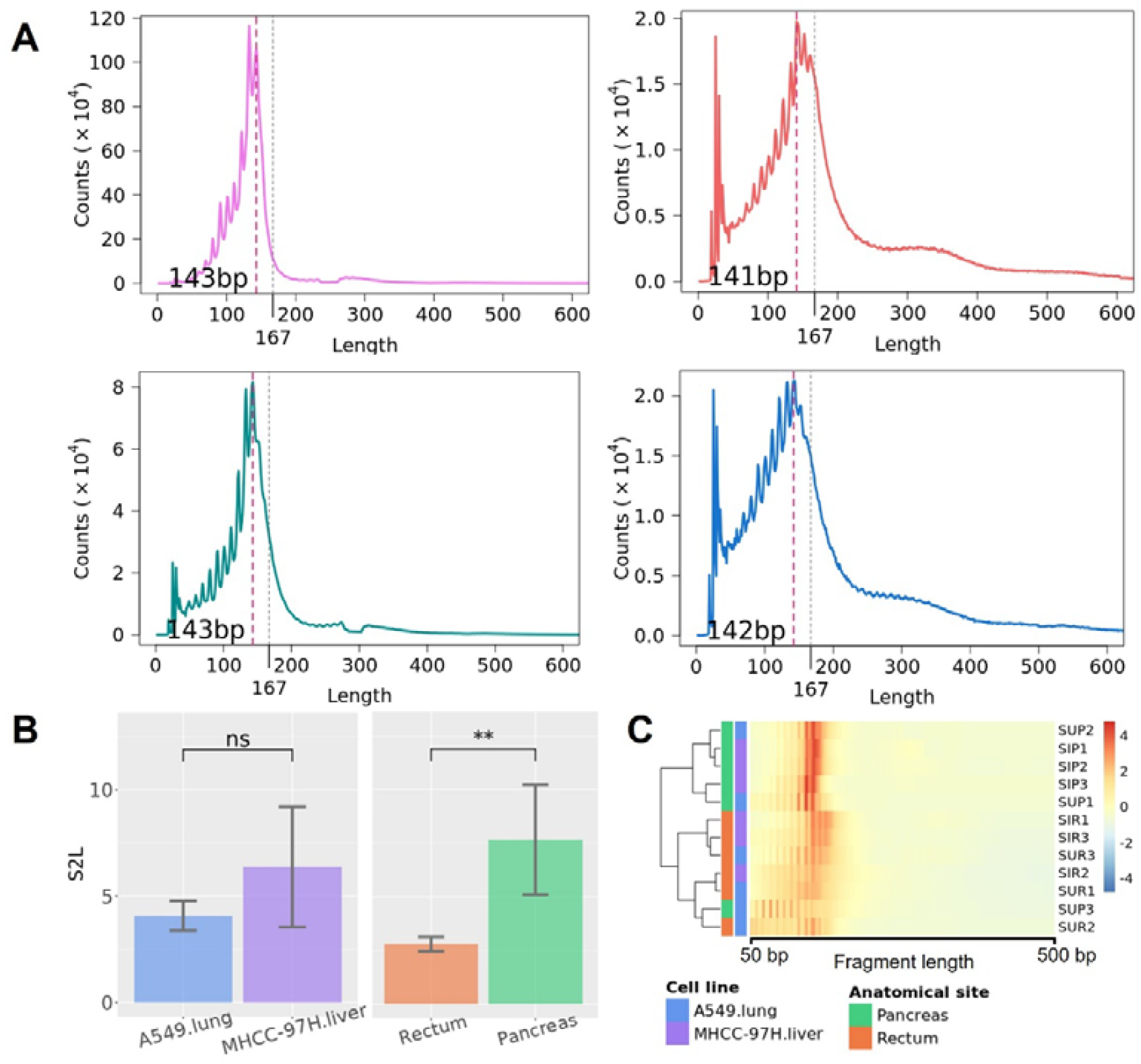
Fragment profile of human tumor cell-derived ctDNA and the enrichment of short fragments. (A) The fragment size distribution of human tumor cell-derived ctDNA from pooled samples representing each of the four types of CDX models, with a major peak around 143 bp. (B) Short ctDNA fragments are significantly more enriched in pancreas CDX models, indicated by S2L ratio. (C) Hierarchical clustering of ctDNA fragment size in CDX models. (significance: ‘*’, p < 0.05; ‘**’, p < 0.01; U-test)

It’s interesting to observe larger variations of ctDNA fragments between anatomical sites than that between cell lines. Distinct fragmentation patterns may result in different fragmentomics features, thus we also calculated BPM, EDM, and FSD for ctDNA fragments. Conversely to CDX-induced cfDNA, samples were mainly clustered by anatomical sites, rather than by cell lines (**Figure 5A S7**). We also observed significantly different AUC distributions when comparing cell lines versus comparing anatomical sites (p < 1e^-4^; U-test) (Figure 5B), resulting in significantly more informative motifs and FSD bins (AUC > 0.9) between pancreas and rectum models than between M97H cell and A549 cell models (p < 0.001; χ2-test) (**Table 1**). The permutation test also demonstrated a significant distinguishment between anatomical sites (p = 0.019) but not between tumor cell lines (p = 0.361). CtDNA from pancreas models included a higher proportion of fragments with sizes between 120-129 bp, while ctDNA from rectum models was sparsely enriched in fragments ranging from 185 bp to 214 bp (**Figure 5C**). These results implied that ctDNA could undergo specific fragmentation in distinct niches during tumorigenesis, resulting in distinguished fragmentomics features likely indicating its anatomical site.

**Figure 5.**
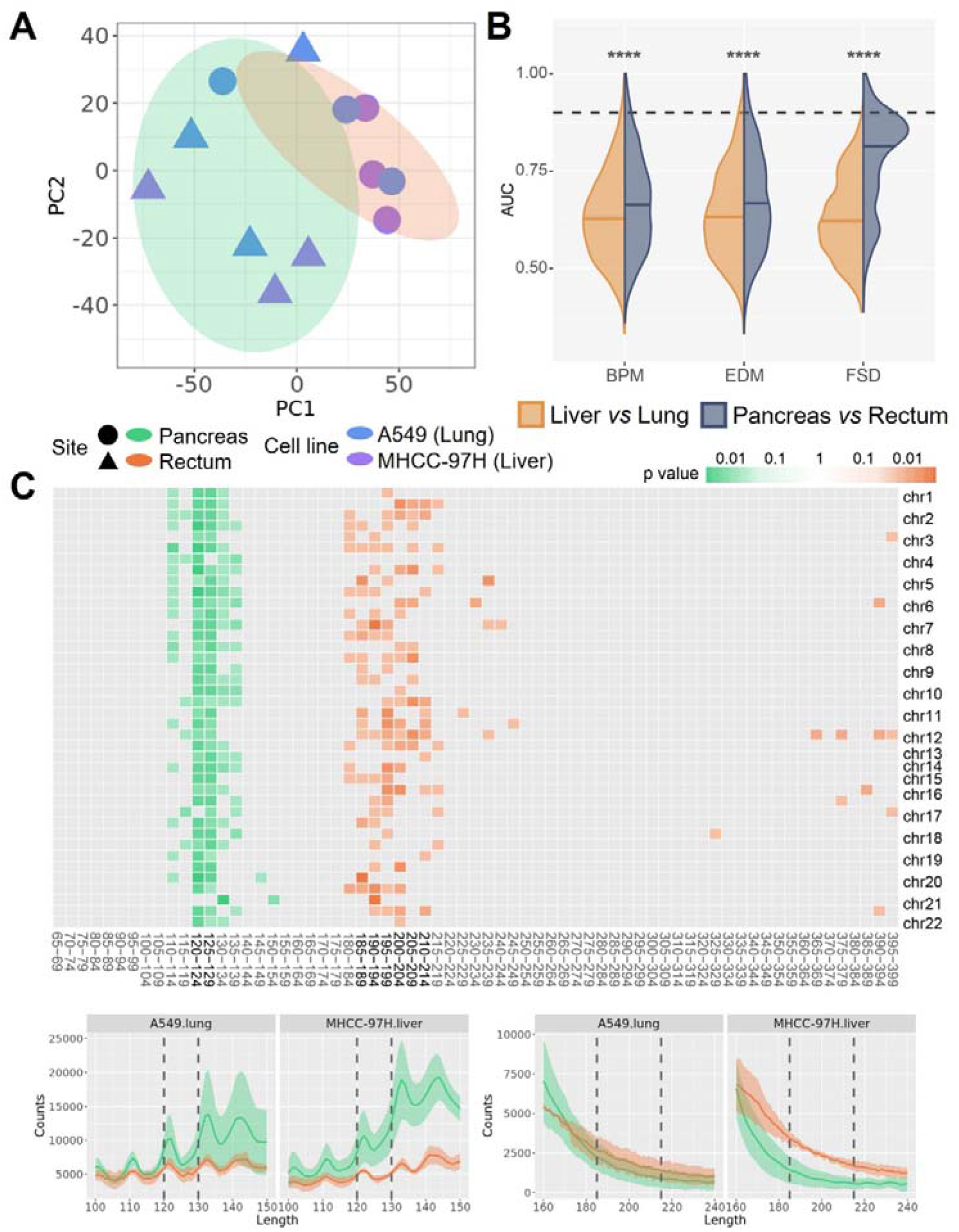
Human tumor cell-derived ctDNA distinguished between CDX models of different anatomical sites. (A) CDX models of different anatomical sites are distinguished by ctDNA fragmentation features like FSD. (B) Comparison of AUC value distributions between samples grouped by tumor cell line and by anatomical site, for fragmentation features including BPM, EDM, and FSD, respectively. The dashed line indicates AUC = 0.9. (significance: ‘****’, p < 1e^-4^; U-test) (C) The fragment bin sizes of FSD that are enriched in pancreas CDX models (green) and rectum CDX models (orange) (top panel), and the differentiation of the fragment profiles corresponding to the bin sizes, with the line indicates the mean value of the replicates and the shadow indicates the standard error (bottom panel). The mosaic was colored if the AUC of the corresponding FSD bin size was larger than 0.9, with the darkness indicating the p value calculated by the U-test.

## Discussion

With the increasing insights into plasma cfDNA biology, it has been actively utilized for clinical non-invasive detection, particularly for early cancer detection. The spectrums of genomic mutations and methylation patterns have been demonstrated to be associated with tumors, and can be easily interrogated from the tumor/para-tumor tissues and/or blood cells, since they are inherited directly from the cellular genome DNA, whereas the fragmentation is an intrinsic feature of cfDNA, which is a mixture of DNA fragments in the bloodstream, with multiple origins. Thus, there is a lack of detailed exploration of the variations attributed to cfDNA fragmentation. Utilizing CDX mouse models, we isolated human-derived ctDNA from CDX-induced cfDNA, and separately investigated their properties, demonstrating that both CDX-induced cfDNA and ctDNA contributed to the variations of cfDNA fragments observed in tumor samples.

Based on the CDX models and control mouse plasma samples, our bioinformatic pipeline generated nearly pure mouse cfDNA fragments and human ctDNA fragments, respectively, minimizing interference with each other. The concentration of ctDNA varied largely among CDX models but had little influence on the differentiation with respect to fragmentation patterns (**Figure S8**). Previous studies have reported an increase in short cfDNA in tumor samples but were unable to tell the source of the elevated short fragments. In the present study, it’s remarkably interesting to observe an enrichment of short fragments in not only human-derived ctDNA but also mouse cfDNA in CDX models. This result suggested that the cells in the tumor niche, regardless of tumor cells, possibly underwent some modifications, which increased the accessibility of DNA and further resulted in short cfDNA fragments from non-tumor cells. Furthermore, these cells were more likely to be hematopoietic cells rather than the ambient tissue cells of the implanting sites; otherwise, we would have observed more significant differences between models of different anatomical sites.

It’s more interesting to observe a larger variation between anatomical sites than that between tumor cell lines in ctDNA fragmentation. Tumor cells implanted in pancreas showed significantly elevated short ctDNA fragments, likely attributable to the aggressiveness of cancers promoted by the tumor microenvironment within the pancreas [39]. Notably, the CNA patterns in ctDNA mainly exhibited the differentiation between tumor cell lines, as expected, while the fragmentation features distinguished the samples between different anatomical sites, implying that fragmentation features might be more variable than genomic alterations. Therefore, they are likely influenced by post-implantation regulation of the tumor cells, or other factors in the tumor microenvironment. Furthermore, we observed converse results that CDX-induced cfDNA and human ctDNA could distinguish samples between different tumor cells and samples between different anatomical sites, respectively, which were confirmed by multiple fragmentation features. Hence, we hypothesized that tumor-induced short cfDNA and ctDNA might stem from different cell types and could be subject to different regulatory mechanisms, producing distinct fragmentation patterns. However, further studies are required to validate the differences between tumor-induced cfDNA and ctDNA, since the sample size was limited in the present study. Other approaches, like methylation sequencing, may be applied to confirm the molecular changes of the DNA fragments, although this is beyond the scope of this study.

Altogether, taking advantage of CDX models, we were able to dissect the fragmentation features of both CDX-induced cfDNA and ctDNA, which was also provided as an ideal resource to study the properties of ctDNA and develop new analytical tools. We demonstrated that both the CDX-induced cfDNA and ctDNA contributed to the fragmentation variations observed in tumor samples, offering a promising metric for cancer prediction. However, caution is warranted in inferring tissue-of-origin, since the fragmentation features may be more variable and susceptible to local regulations in tumor niches.

## Methods

### Establishment of cell line-derived xenograft models

Human MHCC-97H cells (liver carcinoma with high metastatic potential) and A549 cells (lung carcinoma) were cultured in DMEM and DMEM/F-12 medium, respectively, supplemented with 10% fetal bovine serum and 1% penicillin/streptomycin and maintained at 37°C with 5.0% CO_2_.

The cell cultures sustained at suitable concentrations so that 5×10^6^ cells and 2×10^6^ cells were *in situ* injected into pancreas and rectum per mouse, respectively. A total of 12 BALB/C male nude mice, aged 6 weeks, were used for cell line implantation, with 3 replicates for each cell line and each anatomical site for injection. Tumor sizes were monitored twice per week until the mice were sacrificed for blood collection. Mice were deeply anesthetized using Zoletil 50 (VIRBAC Trading (Shanghai) Co., Ltd.), followed by blood collection via cardiac puncture to obtain at least 500 µl of blood, and then humanely euthanized using COD. All experiment procedures were conducted with the approval of the Ethics Committee of the School of Life Sciences at Fudan University.

### Plasma cfDNA extraction and sequencing

Whole blood from the xenograft mice and three normal male BALB/C mice was collected in cfDNA preservation tubes (HiGiA) and immediately stored in 4D until centrifugation. The plasma isolation was processed as previous described [40], using a two-step centrifugation method. The cfDNA (>20 ng) was subsequently extracted from plasma using HiPure Circulating DNA Midi Kit C (Magen). Sequencing library was constructed with all the extracted cfDNA per sample, using KAPA Hyper Prep Kit (Roche), according to the manufacturer’s instructions, during which a 10 bp-UMI was liganded to 3’end of each cfDNA fragment. The Illumina NovaSeq was employed to conduct WGS, with 150 bp paired-end reads.

### Sequencing data processing and ctDNA isolation

The entire WGS sequencing data (fastq) consisted of a mixture of mouse cfDNA fragments and human tumor cell-derived ctDNA fragments. A pipeline was developed to precisely separate the two types of cfDNA fragments. Raw reads were first trimmed by fastp [41] (version 0.20.1). We then employed xenome [36] and XenofilteR [37] to retrieve high-confidence human-derived reads.

The high-quality reads were mapped to the mouse reference genome (mm10) and the human reference genome (hg19) using bwa [42] (version 0.7.5a-r405), respectively, after which XenofilteR was used to extract human-derived reads by comparing human alignments to mouse alignments. The results were subsequently converted into fastq files. Meanwhile, xenome was applied to directly cluster human reads and mouse reads from the trimmed reads, separately. The resulting human-derived reads from both methods were merged and then re-aligned to the human reference genome. Additionally, to build the black regions of the human reference genome, we also attempted to generate human-alignments of the three normal mouse samples using both XenofilteR and xenome. Regions that were supported by either at least two samples or two methods were used to compile the list of black regions. These regions were then utilized to filter the re-alignments of the human-derived reads to generate high-confidence ctDNA fragments. All the alignments were filtered by mapping quality ≥ 30. CfDNA fragments were inferred from the paired-end information from the bam files and included in the analysis if their length fell within the range of 50-1000 bp. Reads from the replicates of the same tumor cell line and the same implanting anatomical site were pooled together to create four representative samples.

### CNA detection and fragmentomics feature calculation

The CNAs of the tumor cell lines were inferred from ctDNA using ichorCNA [43, 44], separately for each CDX model, as well as the four pooled samples. The bin size was set to 1Mb, and the log_2_ ratio of each bin was used to generate the CNA feature matrix, which excluded regions in a blacklist (Duke Excludable Regions) and genomic gap list. Other fragmentomics features, including the BPM, EDM, and FSD, were calculated following a previous study [32]. Briefly, the BPM and EDM were calculated for frequencies of all combinations of 6-bp motif (n=4096) flanking 3bp of both ends and inner ends of each cfDNA fragment, respectively. The FSD was defined as the fraction of fragment sizes falling within different bins ranging from 65 bp to 399 bp, with a step of 5 bp. It was calculated for each chromosome arm of the human genome (except for chr13p, chr14p, chr15p, chr21p, chr22p and chrY). For the mouse plasma cfDNA, only q arms of the mouse genome were included for calculation.

### Permutation and statistical analysis

Permutation was conducted on the CDX models to randomly generate two groups, each consisting of six samples. An AUC value was then calculated for each marker (*e*.*g*., FSD bin) based on the permutated samples, and the number of informative markers (AUC > 0.8) was recorded. The empirical numbers of informative markers from real labels of samples, either by tumor cell line or anatomical site, were compared to the distribution of informative marker numbers derived from 1,000 permutations to calculate the p values.

All the statistical analyses were performed in R. The rank sum test (U-test) or t-test was used to compare two groups of samples as indicated in the main text, with BH correction applied when necessary. AUC values were calculated using ‘pROC’ package (version 1.18.0) in R.

## Ethics approval and consent to participate

The study was approved by the Ethics Committee of the School of Life Sciences at Fudan University.

## Consent for publication

Not applicable.

## Availability of data and materials

The datasets generated have been deposited into the Genome Sequence Archive for Human (GSA-Human) of China National Genomics Data Center (NGDC) under accession number PRJCA026522.

## Authors’ contributions

H.S. and D.L. conceived and designed the study. H.S., Y.T. and H.C. performed the experiments. R.F. and H.S. performed data analysis and interpretation. R.F., H.S., and D.L. wrote the manuscript. D.L. provided project supervision. All authors contributed to manuscript editing and revision. All authors approved the submission of the final manuscript for publication.

## Acknowledgements

The authors thank Hua Chen, Chengcheng Ma, Minjie Xu, Wei Li, Chengxiang Gong and Zhixi Su for helpful discussions.

