## Supplementary figures for "Dissecting cell-free DNA fragmentation variation in tumors using cell line-derived xenograft mouse"

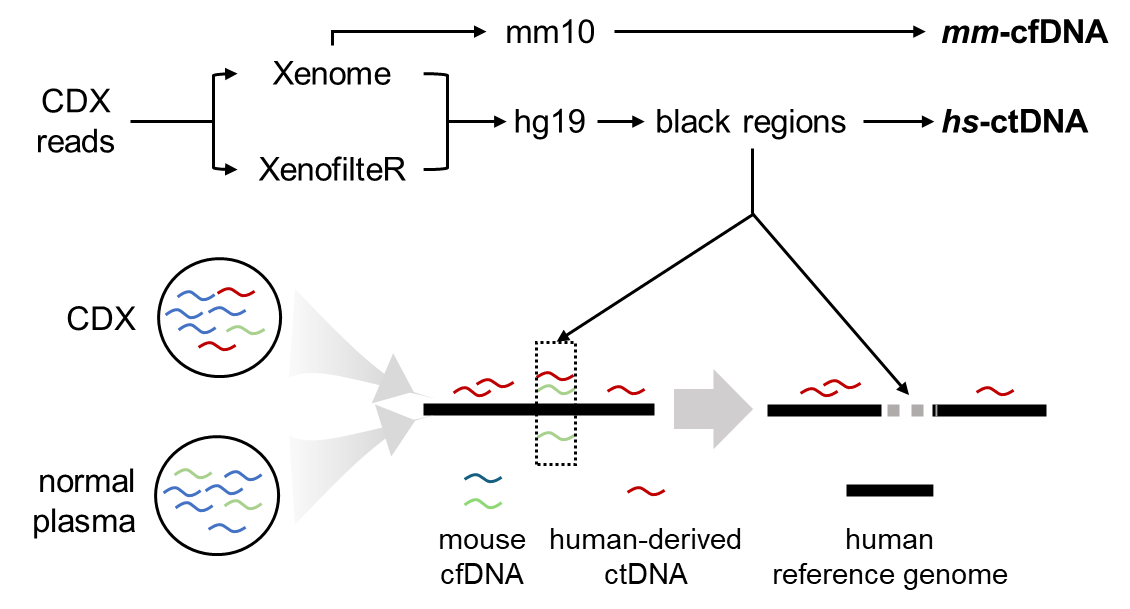

**Figure S1. A bioinformatic pipeline to isolate pure human-derived ctDNA from the xenograft mouse model, including the establishment of a list of black regions.**

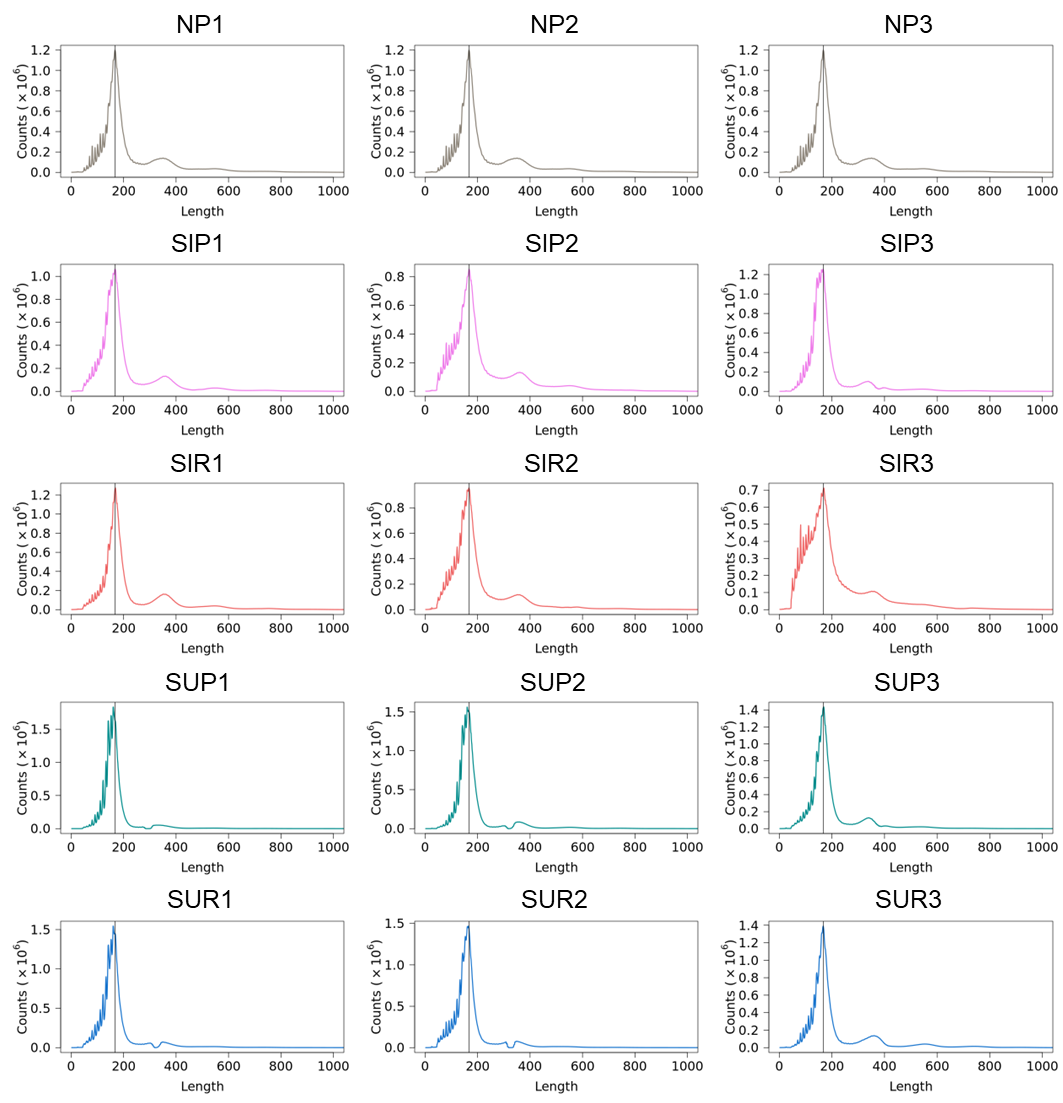

**Figure S2. Fragment size distribution of individual mouse plasma cfDNA, with a mode size of ~167 bp.**

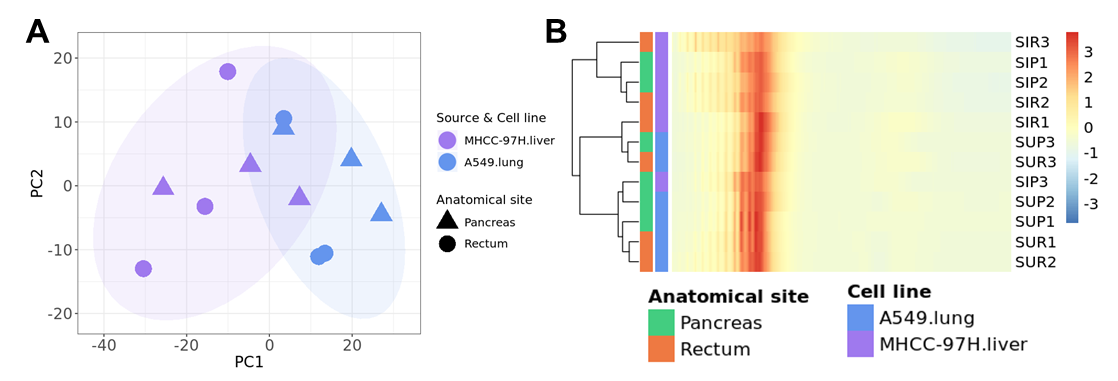

**Figure S3. Principal component analysis (PCA) (A) and hierarchical clustering (B) of the fragment sizes of xenograft mouse plasma cfDNA.**

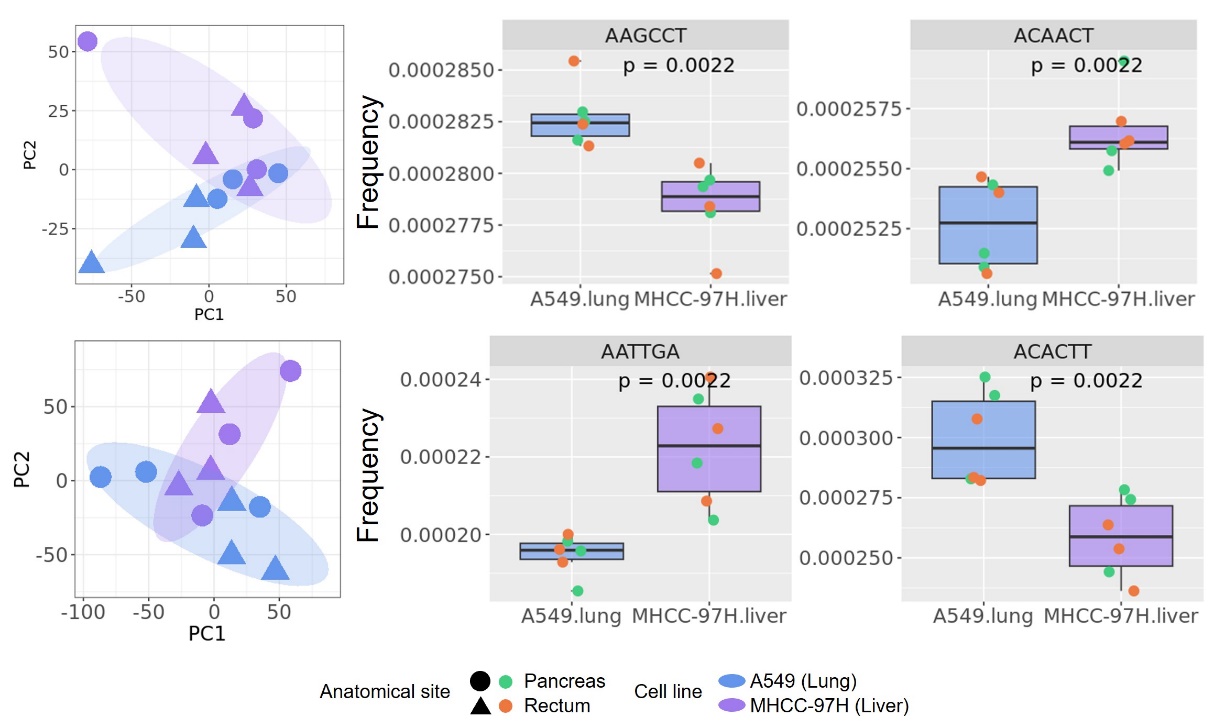

**Figure S4. CDX models of different cell lines are distinguished mainly by CDX-induced cfDNA fragmentation feature of breakpoint motif (BPM) and end motif (EDM), with several cases of the significantly differentiated motifs shown. P values calculated by U-test.**

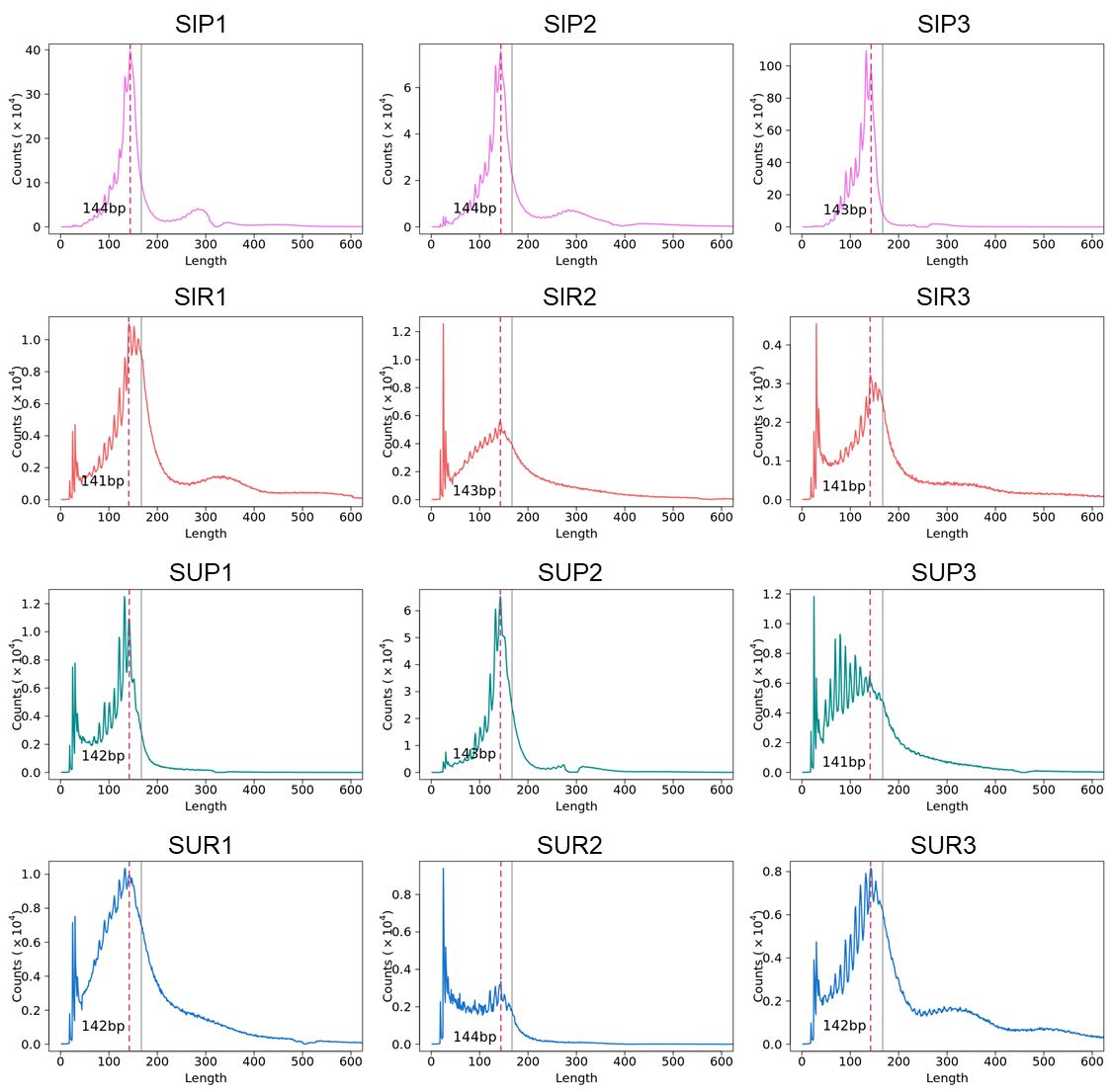

**Figure S5. Fragment size distribution of individual ctDNA from CDX models, with a mode size of ~143 bp.**

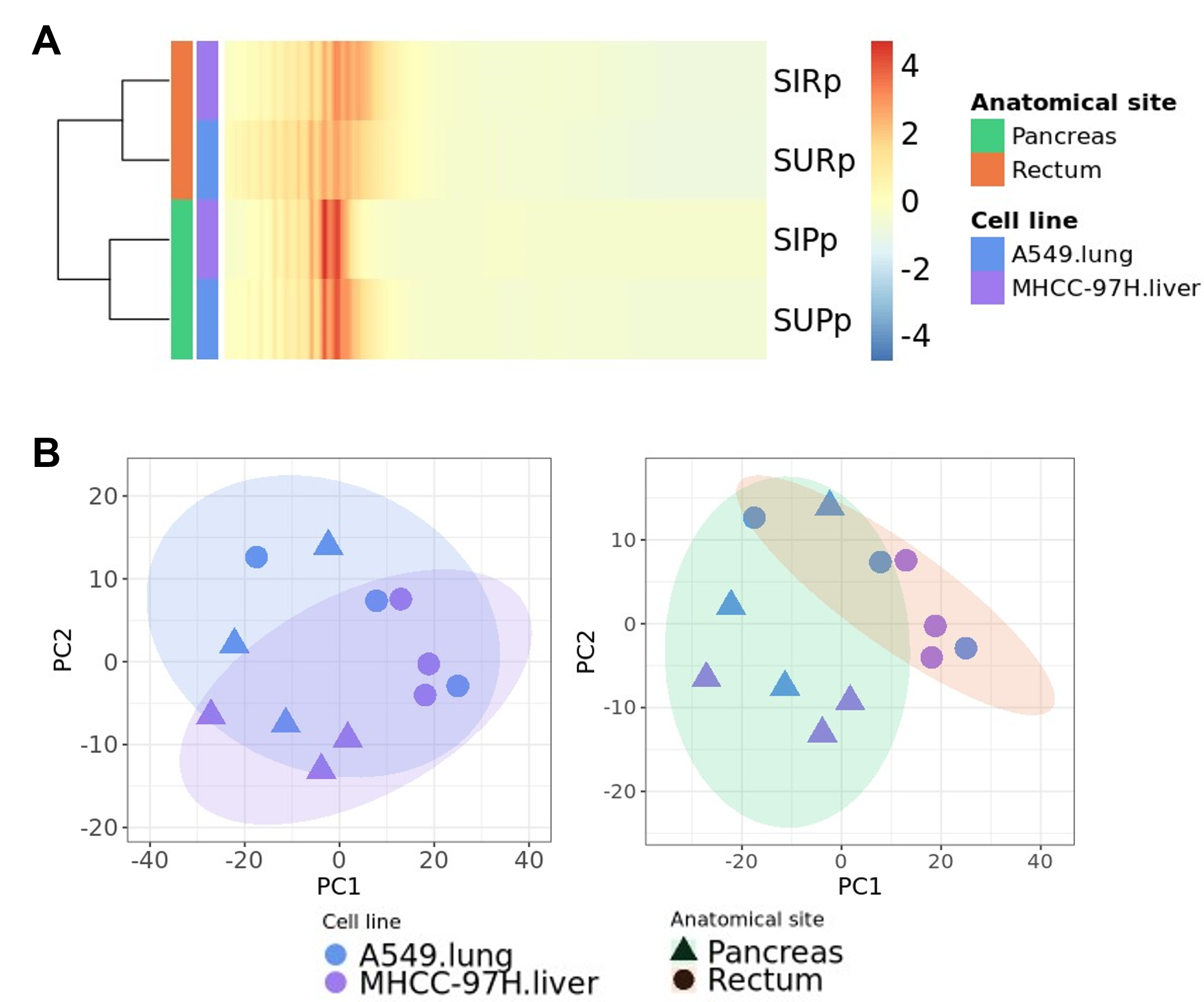

**Figure S6. (A) Hierarchical clustering by the fragment sizes of ctDNA from pooled representative samples. (B) PCA by fragment sizes of ctDNA from CDX models, grouped by different cell lines (left) and by different anatomical sites (right).**

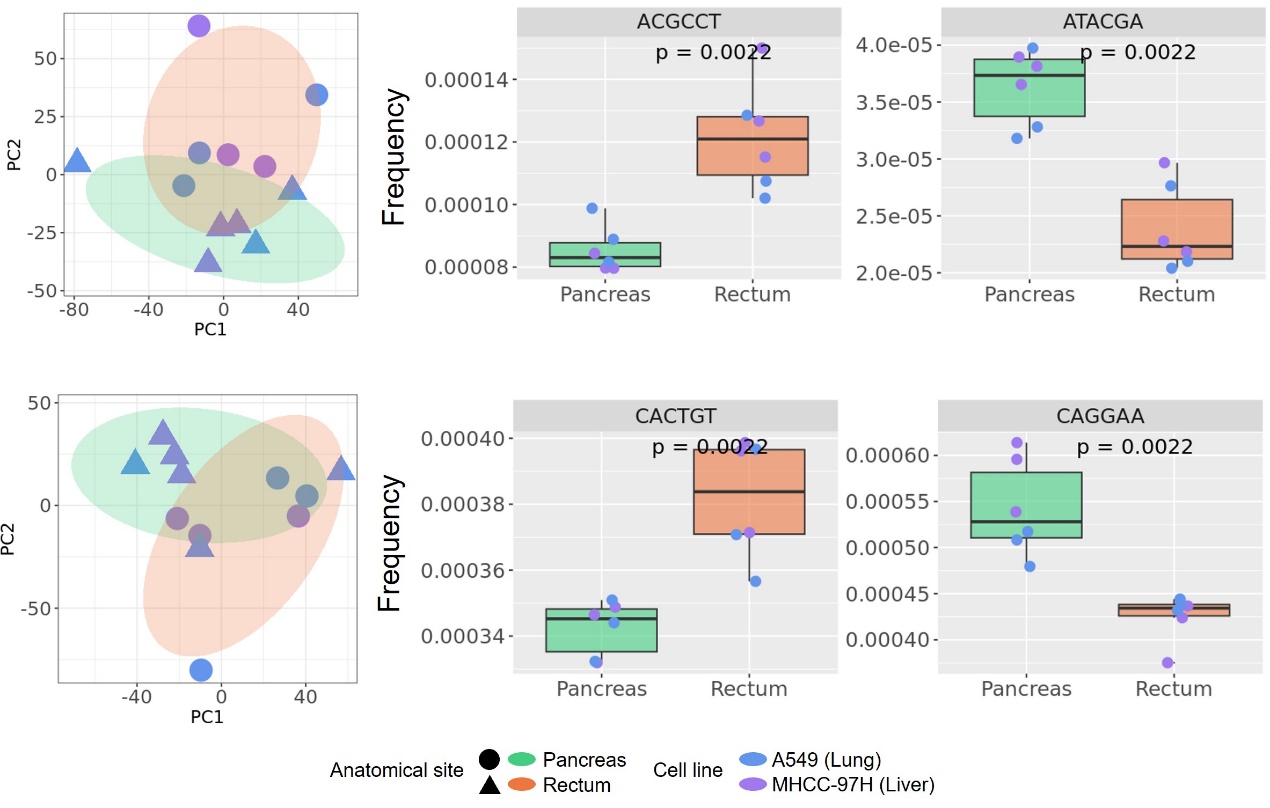

**Figure S7. CDX models of different anatomical sites are distinguished mainly by ctDNA fragmentation feature of BPM and EDM, with several cases of the significantly differentiated motifs shown. P values calculated by U-test.**

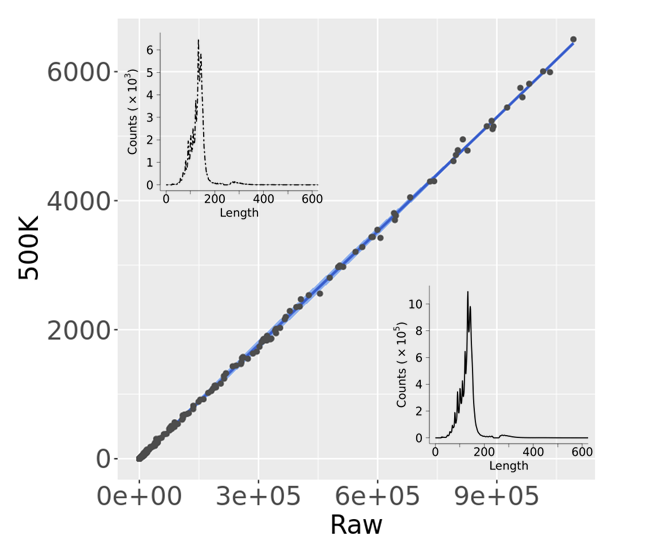

**Figure S8. Correlation of fragmentation profiles between raw ctDNA data and down-sampled ctDNA data (~500K reads), with the inset plots showing the corresponding fragment size distributions.**
